# Cortical and subcortical neurons discriminate sounds in noise on the sole basis of acoustic amplitude modulations

**DOI:** 10.1101/528653

**Authors:** S. Souffi, C. Lorenzi, C. Huetz, J-M Edeline

## Abstract

Humans and animals maintain accurate sound discrimination in the presence of loud sources of background noise. It is commonly assumed that this ability relies on the robustness of auditory cortex responses. However, no attempt has been made to characterize neural discrimination of sounds masked by noise at each stage of the auditory system and disentangle the sub-effects of noise, namely the distortion of temporal cues conveyed by modulations in instantaneous amplitude and frequency, and the introduction of randomness (stochastic fluctuations in amplitude). Here, we measured neural discrimination between communication sounds masked by steady noise in the cochlear nucleus, inferior colliculus, auditory thalamus, primary and secondary auditory cortex at several signal-to-noise ratios. Sound discrimination by neuronal populations markedly decreased in each auditory structure, but collicular and thalamic populations showed better performance than cortical populations at each signal-to-noise ratio. Comparison with neural responses to tone-vocoded sounds revealed that the reduction in neural discrimination caused by noise was mainly driven by the attenuation of slow amplitude modulation cues, with the exception of the cochlear nucleus that showed a dramatic drop in discrimination caused by the randomness of noise. These results shed new light on the specific contributions of subcortical structures to robust sound encoding, and demonstrate that neural discrimination in the presence of background noise is mainly determined by the distortion of the slow temporal cues conveyed by communication sounds.

## Introduction

Understanding the neural mechanisms used by the central auditory system to extract and represent relevant information for discriminating communication sounds in a variety of acoustic environments is a major goal of auditory neurosciences. This enterprise is motivated by the repeated observation that humans and animals successfully maintain high discrimination performance for speech and behaviorally salient calls when the latter are embedded into loud sources of background noise produced by environmental medium (such as tropical forests, underwater or urban environments)^1-6^.

Previous studies have assumed that this perceptual robustness mainly relies on the capacity of cortical neurons to extract invariant features^5,7-10^. For example, in the cortical field L (the analogous of primary auditory cortex (A1) in bird), the percentage of correct neuronal discrimination between zebra-finch songs embedded in different types of acoustic maskers decreases proportionally to the target-to-masker ratio and parallels behavioral performance^5^. Similarly, in a secondary auditory area, neurons generate background-invariant representation of vocalizations at signal-to-noise ratios that match behavioral recognition thresholds^10^. More recently, between-vowels discrimination performance of neuronal populations located in A1 was found to resist to a large range of acoustic alterations (including changes in fundamental frequency, spatial location, or level) and was similar to behavioral performance^9^.

The goal of the present study was to identify the auditory brain structures responsible for these robust neural computations and clarify the effects of background noise on the neural representation of communication sounds, knowing that external noise has three disruptive sub-effects on communication sounds^11,12^: noise attenuates the power of their amplitude modulation cues (AM, also called “temporal-envelope”)^13-15^, corrupts their frequency modulation cues (FM, also called “temporal fine structure”)^16,17^, and introduces stochastic fluctuations in level, that is statistical variability of the AM power^14^. To address these issues, we investigated whether the ability of populations of auditory neurons to discriminate between communication sounds belonging to the same category (e.g. the alarm call in guinea pig) and masked by external noise *increases or decreases* along the auditory pathway from the first auditory relay (the cochlear nucleus) up to the primary and secondary cortical areas. An increased ability may result from the specialization of cortical responses for detecting crucial vocalization features^10,18,19^, whereas a decreased ability may result from the loss of spectro-temporal details promoting the identification of auditory objects^20, 21^. For the first time, the discrimination performance of neuronal populations recorded along the auditory pathway, from the cochlear nucleus up to a secondary auditory cortex, was assessed for four utterances of the same vocalization presented against a stationary broadband noise using three signal-to-noise ratios (SNRs). The results were compared to the effects of an artificial signal-processing scheme (a tone vocoder) that progressively degraded acoustic AM and FM cues (within 38 to 10 frequency bands) without introducing any stochastic fluctuation as in the case of background noise. AM and FM spectra of communication sounds were computed at the output of simulated cochlear filterbank for each acoustic alteration allowing us to identify masking noise and vocoder conditions matched in terms of amount of modulation reduction. We then correlated these reductions of temporal modulation cues with the discriminative neuronal performance recorded in each structure. We demonstrate that, with the noticeable exception of the cochlear nucleus, the larger the reduction of AM cues (the first sub-effect of background noise), the larger the decrease in discriminative abilities in cortical and subcortical structures. Corruption of FM cues (the second sub-effect of noise) had little observable effects if any on neural discrimination. Introduction of stochastic fluctuations (the third sub-effect of noise) impacted neural discrimination in the cochlear nucleus only. In addition, this study revealed that, for each acoustic distortion tested here, the highest level of discrimination was found in subcortical structures, either at the collicular level (in masking-noise conditions) or at the thalamic level (in vocoder conditions).

## Results

From a database of 2334 recordings collected in the different auditory structures, two criteria were used to include neuronal recordings in our analyses. A recording had to show a significant STRF (see Methods) and an evoked firing rate significantly above spontaneous firing rate (200 ms before each original vocalization) for at least one of the four original vocalizations. Applying these two criteria led to the inclusion of 499 recordings in CN, 386 recordings in CNIC, 262 recordings in MGv, 354 recordings in A1 and 95 recordings in VRB (see supplementary Table 1). In the following sections, the neuronal responses to the original vocalizations (see Fig. 1a) presented in quiet are compared across brain structures and the discriminative abilities are described at the individual and population level. The neuronal discriminative abilities tested at the cortical and subcortical level with tone vocoded vocalizations (Fig. 1b) and vocalizations presented against different levels of masking noise (Fig. 1c) are described and compared next.

**Figure 1.**
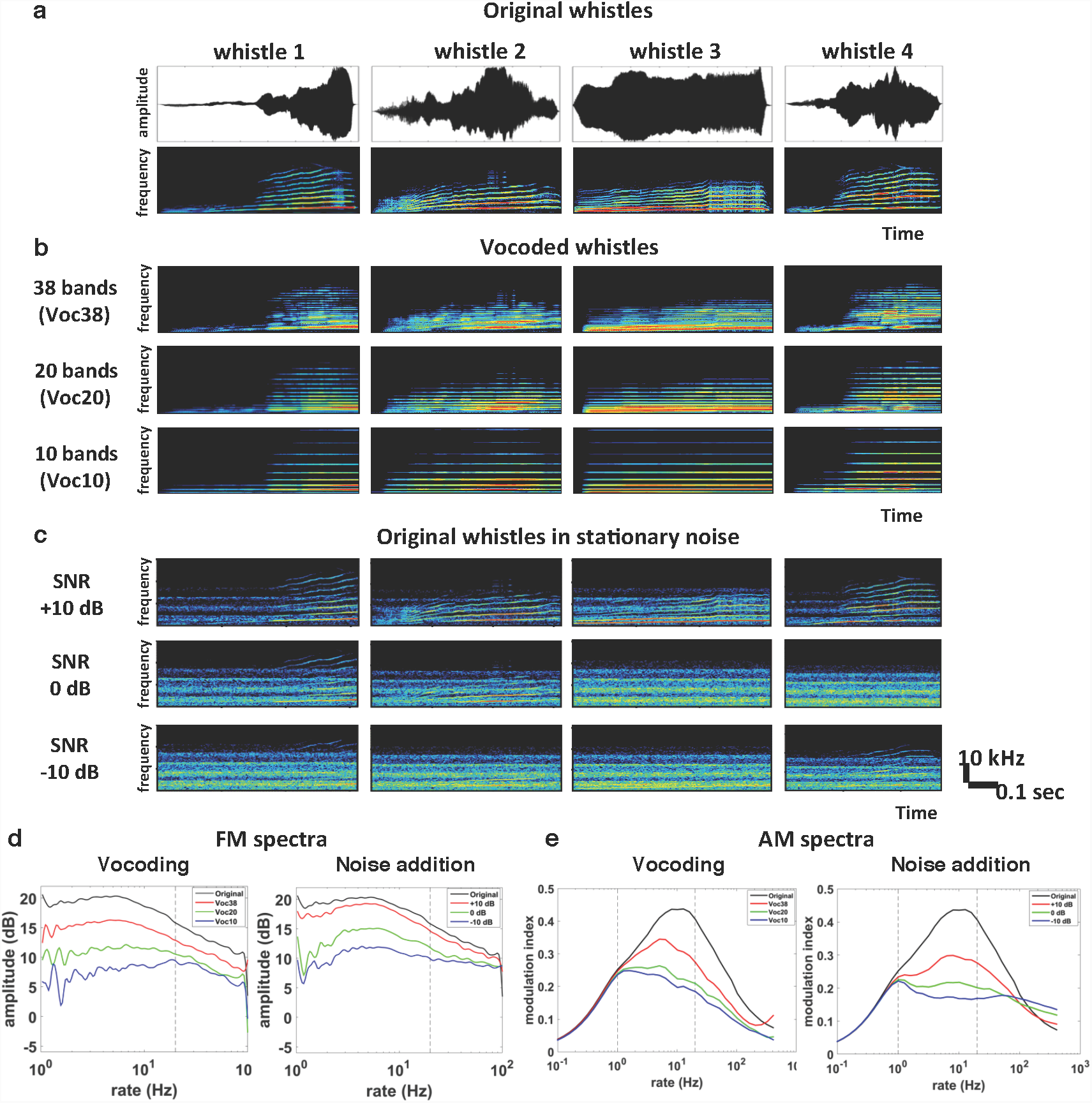
Acoustic stimuli and averaged modulation spectra. **a.** Waveforms (*top*) and spectrograms (*bottom*) of the four original whistles used in this study. **b.** F*rom top to bottom,* spectrograms of the four vocoded whistles using 38, 20 and 10 frequency bands. **c.** F*rom top to bottom, s*pectrograms of the four original whistles embedded in stationary noise at three SNRs (*+10, 0* and *-10 dB*) and spectrograms of the stationary noise only (*Noise only*). **d.** Vocoding and noise effects on frequency-modulation (FM) spectra. The two plots represent the averaged modulation spectra of the four original vocalizations (*in black*), vocoded vocalizations (*Voc38, Voc20* and *Voc10: red, green and blue respectively, left panel*) and vocalizations in stationary noise at three SNRs (*+10, 0* and *-10 dB* : *red, green and blue respectively, right panel)* **e.** Vocoding and noise effects on amplitude-modulation (AM) spectra. The two plots represent the averaged modulation spectra of the four original vocalizations (*in black*), vocoded vocalizations (*Voc38, Voc20* and *Voc10: red, green and blue respectively, left panel*) and vocalizations in stationary noise at three SNRs (*+10, 0* and *-10 dB* : *red, green and blue respectively, right panel)*. Vertical black dashed lines on AM and FM spectra correspond to the frequency range (1-20 Hz) selected for the data analysis.

### Discrimination of the original vocalizations in quiet culminates at the subcortical level

Figure 2a displays neuronal responses of two simultaneous recordings obtained at five levels of the auditory pathway (CN, CNIC, MGv, AI and VRB). The neuronal responses were strong and sustained in the three subcortical structures whereas they were more phasic in AI and more diffuse in VRB. For most of the recordings, temporal patterns of responses were clearly reproducible from trial-to-trial, but they differed from one vocalization to another both at the cortical and subcortical level.

**Figure 2.**
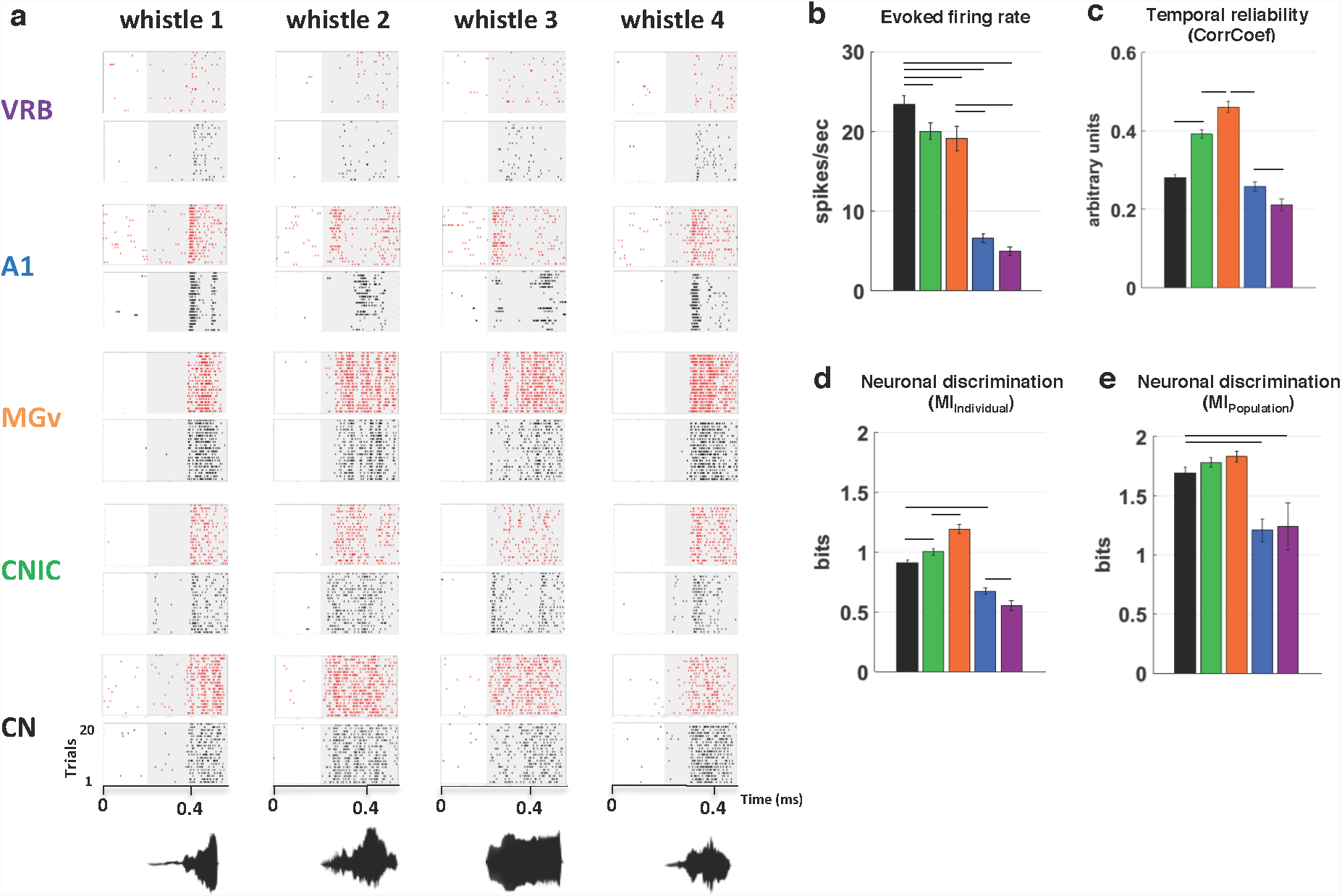
Subcortical neurons discriminate better the original vocalizations than cortical neurons. **a.** *From bottom to top, n*euronal responses were recorded in CN, CNIC, MGv, A1 and VRB simultaneously under 16 electrodes but only two are represented here, with alternated black and red colors. Each dot represents the emission of an action potential and each line corresponds to the neuronal discharges to one of four original whistles. The grey part of rasters corresponds to evoked activity. The waveforms of the four original whistles are displayed under the rasters. **b-e.** The panels show (**b**) the evoked firing rate (spikes/sec), (**c**) the temporal reliability quantified by the CorrCoef value (arbitrary units), (**d)** the neuronal discrimination assessed by the mutual information (MI) computed at the level of the individual recording (MI_Individual_, bits) and (**e**) at the level of neuronal population (MI_Population_, bits) with populations of 9 simultaneous recordings obtained with the four original vocalizations in CN (*in black*), CNIC (*in green*), MGv (*in orange*), A1 (*in blue)* and VRB *(in purple).* In each structure, error bars represent the SD of the mean values and black lines represent significant differences between the mean values (unpaired *t* test, p<0.05). Note that the evoked firing rate decreases from the CN to VRB but both the temporal reliability (CorrCoef) and the discriminative ability (MI) values reach a maximal value in MGv. Note also that at the population level, all the subcortical structures discriminate better the original vocalizations than cortical areas.

Quantifications of evoked responses to original vocalizations are presented on Figures 2b-e for each auditory structure. These analyses clearly pointed out large differences between the mean values of evoked firing rate, CorrCoef and MI quantified at the cortical vs. at the subcortical level. First, the evoked firing rate was significantly higher in the subcortical structures than in the cortex (unpaired t-test, lowest p value p<0.001). It was also higher in CN than in the other subcortical structures (Fig. 2b). Second, the CorrCoef values were significantly higher in the subcortical structures than in AI and VRB (Fig. 2c), indicating that the trial-to-trial reliability of evoked responses was stronger at the subcortical than at the cortical level, reaching its maximum in the CNIC and MGv. CorrCoef value significantly increased from CN to CNIC (unpaired t-test, p<0.01), and then from CNIC to MGv (p<0.01; Fig. 2c). CorrCoef decreased significantly between the two cortical areas and CNIC and MGv (p<0.01); it also decreased relative to CN (p=0.09). The CorrCoef value was also significantly lower in VRB than in AI (p=0.035). Third, the MI_Individual_ values found at the subcortical level were significantly higher than at the cortical level (unpaired t-test, highest p<0.001 between the cortex and the other structures; Fig. 2d). At the subcortical level, the MI_Individual_ value was significantly higher in MGv than in CNIC and CN (unpaired t-test, p<0.01). The MI_Individual_ value was also significantly lower in VRB than in AI (p = 0.037). The highest mean MI_Individual_ value was found in MGv, suggesting that, on average, thalamic neurons discriminate better the four original whistles than the other auditory structures. Note that this high MI_Individual_ value is related to the higher temporal trial-to-trial reliability (indexed by the CorrCoef value) obtained at the thalamic level (see Fig. 2c).

The distributions of MI_Individual_ value were plotted as a function of temporal precision for each structure (see supplementary Fig. 2) to investigate whether the higher mean MI_Individual_ values found in subcortical responses result from the fact that neurons in these structures systematically conveyed more information about the stimuli than cortical neurons. With a temporal resolution of 8 ms, the proportion of neurons having a MI_Individual_ curve reaching a value of 1.5 bits (indicating that at least 3 stimuli can be discriminated) was similar in CN and CNIC (22% and 21% respectively, chi-square test <1; p=0.99); it was significantly higher in MGv (39%, p=0.017 and p=0.04) and lower in AI (3.5%, p=0.001 in both cases) and in VRB (2%, p=0.001).

Finally, MI was also computed based on the temporal patterns obtained from two to sixteen simultaneous recordings to determine whether the discriminative abilities of neural networks confirm the results obtained at the individual (i.e., single unit) level. MI_Population_ quantifies how well the four whistles can be discriminated based on temporal patterns expressed by populations distributed on the tonotopic map. Figure 2e presents the MI_Population_ computed from 9 simultaneous recordings for the five structures under investigation: This figure confirms that neural populations in subcortical structures discriminate the four original whistles better than the cortical populations (unpaired t-test, highest p value p<0.002 between CN and VRB) without any statistical difference between the three subcortical structures. An examination of the evolution of the MI_Population_ as a function of the number of simultaneous recordings in the different structures revealed that the growth functions rapidly reached high values in all subcortical structures, whereas there were only a few of such curves in AI and VRB whatever the number of recordings considered (see supplementary Fig. 3).

### Modest effects of tone vocoding

Figure 3a displays rasters of recordings obtained in the five structures in response to the original and tone vocoded vocalizations. As illustrated here, at all levels, neurons still responded to the vocoded stimuli even for 10 frequency-band vocoded stimuli. However, the firing rate was decreased in each structure, and so was the precise organization of neuronal responses, especially with the 10 frequency-band vocoded stimuli.

**Figure 3.**
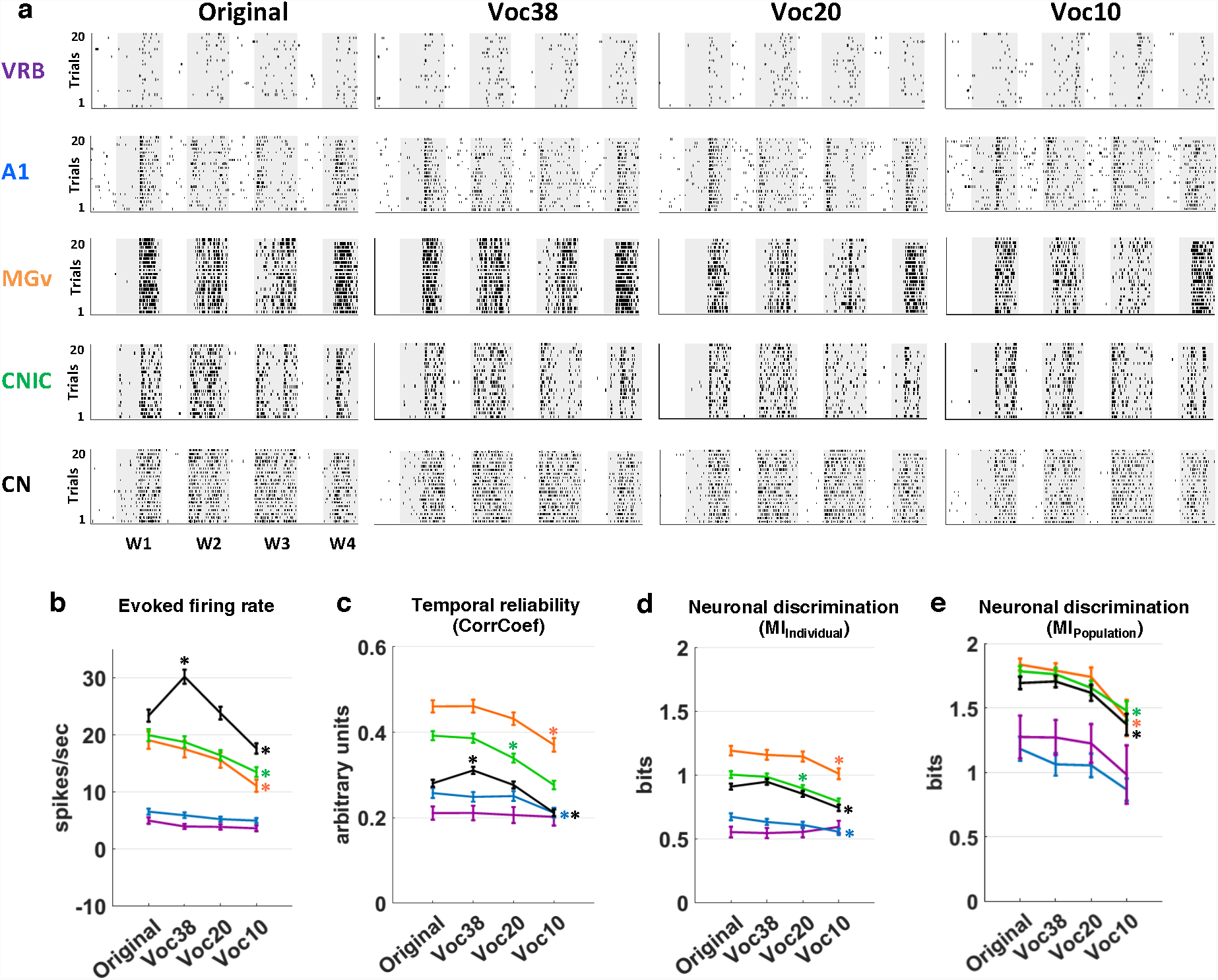
Vocoding slightly alters neuronal responses at each stage of the auditory system. **a.** *From left to right*, raster plots of responses to the four original whistles (*Original*) and their vocoded versions generated using either 38, 20 or 10 frequency bands (*Voc38, Voc20 and Voc10). From bottom to top,* neuronal responses were recorded in CN, CNIC, MGv, A1 and VRB. **b-e.** The mean values (±SEM) represent (**b**) the evoked firing rate (spikes/sec), (**c**) the temporal reliability represented by the CorrCoef value (arbitrary units), (**d**) the neuronal discrimination assessed by the mutual information (MI) computed at the level of the individual recordings (MI_Individual_, bits) and (**e**) at the level of neuronal population (MI_Population_, bits) with populations of 9 simultaneous recordings obtained with original (*Original*) and vocoded vocalizations (*Voc38, Voc20 and Voc10)* in CN (*in black*), CNIC (*in green*), MGv (*in orange*), A1 (*in blue)* and VRB *(in purple) (one-way ANOVA,* *P < 0.05). At the population level, the discriminative abilities significantly decreased only for 10 frequency bands in subcortical structures and did not decrease in cortical areas.

Figures 3b-e summarize vocoding effects on the four parameters quantifying neuronal responses. Apart from an initial increase in firing rate observed only in CN with the 38-band vocoded stimuli, the effects on evoked firing rate were modest in each structure (Fig. 2b): A significant decrease in evoked firing rate between the responses to the original and the 10-band vocoded vocalizations was only found at the subcortical level (for all subcortical structures, ANOVA test: p<0.001, F_CN(3,1995)_=22.6; F_CNIC(3,1543)_=8.85; F_MGv(3,1047)_=6.55), whereas there was no decrease in either AI or VRB. Vocoding also decreased the mean CorrCoef values in each structure except in VRB (Fig. 3c): this decrease was significant with the 10-band vocoded vocalizations in CN, MGv and in AI (ANOVA test, highest p value, p<0.02, F_CN(3,1930)_=26.48, F_MGv(3,889)_=7.7, F_A1(3,1125)_=3.42). The decrease in CorrCoef value was already significant with 20-band vocoded vocalizations in the CNIC (p<0.001, F_(3,1391)_=26.19).

Similarly, tone vocoding decreased the MI_Individual_ values in each structure except in VRB (Fig. 3d). Here too, the decrease was significant with the 10-band vocoded vocalizations in CN, MGv and AI (ANOVA test, highest p value, p<0.02, F_CN(3,1445)_=12.23, F_MGv(3,810)_=3.75, F_A1(3,720)_=3.59) and it was already significant with 20-band vocoded vocalizations in the CNIC (p<0.001, F_(3,1231)_=13.17). At the population level, there was a striking difference between the subcortical and cortical structures (Fig. 3e): compared with the values obtained with original vocalizations, the MI_Population_ values computed with the 10-band vocoded vocalizations were significantly lower in the subcortical structures (ANOVA test, highest p value, p<0.005, F_CN(3,127)_=6.46, F_MGv(3,67)_=4.62, F_CNIC(3,115)_=6.28) but not at the cortical level. The evolution of MI_Population_ as a function of the number of simultaneous recordings (see supplementary Fig. 4a) indicated that in each subcortical structure, the curves rapidly reached high MI_Population_ values (close to the maximal value of 2) in each vocoding conditions, whereas there were only a few of such curves in AI and VRB whatever the vocoding condition.

In conclusion, for the five auditory structures under study, the neuronal responses to 10-band vocoded vocalizations were slightly weaker, temporally less accurate and less discriminative than the responses to the original vocalizations. Nonetheless, subcortical neurons still maintained the highest ability to discriminate between tone vocoded vocalizations, both at the level of individual recordings and at the population level.

### Pronounced effects of masking noise on neuronal discrimination

The rasters presented in figure 4a show the effects produced by presenting the original vocalizations against a stationary noise at three SNRs (+10, 0 and −10 dB). Masking noise attenuated neuronal responses at each level of the auditory system. However, auditory structures were differentially affected by noise. In these rasters, the responses in the CNIC did not change up to a 0 dB SNR, decreasing only at a −10 dB SNR. This was not the case in the other auditory structures where the responses decreased either at a +10 dB SNR (MGv and CN) or at a 0 dB SNR (AI and VRB).

**Figure 4.**
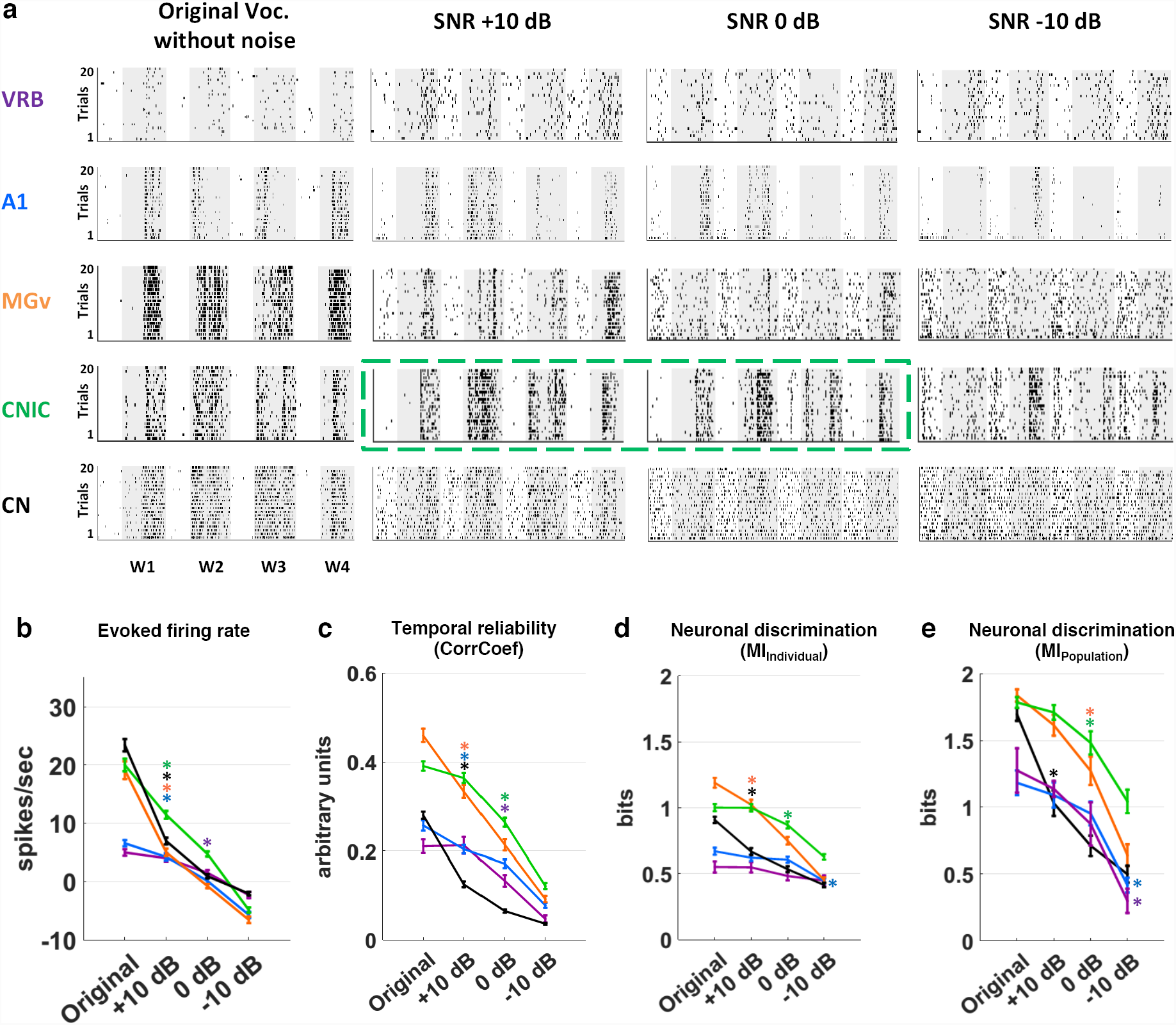
Noise strongly reduces neuronal responses in all structures but to a lesser extent in the central nucleus of the inferior colliculus. **a.** *From left to right*, raster plots of responses of four original whistles (*Original*) and their noisy versions in stationary noise at three SNRs: +10, 0 and −10 dB. *From bottom to top,* neuronal responses were recorded in CN, CNIC, MGv, A1 and VRB. The grey area corresponds to the evoked activity for each vocalization. The green dashed lines show a typical example of CNIC neuronal responses that are resistant to the noise addition. **b-e**. The mean values (±SEM) represent (**b**) the evoked firing rate (spikes/sec), (**c**) the temporal reliability represented by the CorrCoef value (arbitrary units), (**d**) the neuronal discrimination assessed by the mutual information (MI) computed at the level of the individual recordings (MI_Individual_, bits) and (**e**) at the level of neuronal population (MI_Population_, bits) with populations of 9 simultaneous recordings obtained with original and vocalizations in stationary noise at three SNRs (+10, 0 and −10 dB SPL) in CN (*in black*), CNIC (*in green*), MGv (*in orange*), A1 (*in blue)* and VRB *(in purple) (one-way ANOVA,* *P < 0.05). Note that at the population level, the discriminative abilities significantly decreased in all structures when SNR decreased, with the CNIC populations still able to discriminate 2 out of 4 stimuli (MI_Population_ value >1).

Figures 4b-e summarize the effects of masking noise on the different parameters quantifying neuronal responses. Masking noise significantly reduced the evoked firing rate in each auditory structure as early as the +10 dB SNR (Fig. 4b, ANOVA test: p<0.001, F_CN(3,1995)_=309.33, F_CNIC(3,1543)_=220.64, F_MGv(3,1047)_=155.07, F_A1(3,1415)_=96.27), except in VRB. Masking noise strongly reduced the CorrCoef values in CN and MGv at the highest (+10 dB) SNR tested here (Fig. 4c; ANOVA test, p<0.001, F_CN(3,1884)_=382.22, F_MGv(3,791)_=155.82) whereas in the CNIC, this reduction was significant only at the 0 dB SNR (ANOVA test, p<0.001, F_(3,1357)_=154.12). At the cortical level, the CorrCoef values were significantly reduced in AI at the +10 dB SNR and in VRB at the 0 dB SNR (ANOVA test, p<0.001, F_A1(3,1093)_=60.83, F_VRB(3,335)_=29.56). At the cortical level, noise significantly reduced the mean MI_Individual_ value in AI at the −10 dB SNR (ANOVA test, p<0.001, F_(3,649)_=9.49) whereas the mean MI_Individual_ value in VRB remained unchanged. At the subcortical level, noise reduced the MI_Individual_ values but again, there was a marked difference between the CNIC and the other subcortical structures: the mean MI_Individual_ value in CN and MGv was significantly reduced at the +10 dB SNR (Fig. 4d; ANOVA test, p<0.001, F_CN(3,819)_=56.75, F_MGv(3,621)_=63.61), whereas the MI_Individual_ value in the CNIC was only significantly reduced at the SNR of 0 dB (ANOVA test, p<0.001, F_(3,1078)_=32.08). Note, however, that at least 20% of the CN recordings maintained MI_Individual_ values above 1 bit, suggesting that a sub-population of CN neurons still sent information about the vocalization identity at higher brainstem centers (see Supplementary Fig. 5). This specific sub-population of CN neurons did not display parameters of their STRFs quantification that differ from the neurons exhibiting MI_Individual_ values below 1 bit at the +10 dB SNR.

In noise conditions, MI_Population_ also allowed to characterize the effects of masking noise on the network discriminative abilities (Fig. 4e). At the cortical level, there was a significant reduction of MI_Population_ values only at the −10 dB SNR (ANOVA test, p<0.001, F_A1(3,111)_=16.63, F_VRB(3,23)_=11.41) whereas there was a significant decrease in CN as early as the +10 dB SNR (ANOVA test, p<0.001, F_(3,127)_=51.49). In MGv and CNIC, neuronal populations displayed the highest discriminative abilities although the decrease in MI_Population_ value was significant at the 0 dB SNR (ANOVA test, p<0.001, F_MGv(3,67)_=41.59, FCNIC(3,115)=22.59).

The evolution of the MI_Population_ as a function of the number of simultaneous recordings in the different structures (see Supplementary Fig. 4b) revealed that whatever the number of neurons considered, noise effects were similar: the population curves in CNIC and MGv grew up relatively rapidly and reached higher values than the curves obtained in CN and in the two cortical areas whatever the SNR.

To summarize, masking noise reduced similarly firing rate in each structure but impacted differently the neurons’ discriminative abilities. Although cortical neurons were the most resistant to changes in noise level, the thalamic and collicular neurons maintained higher MI values, with the CNIC neurons displaying the highest discriminative abilities both at the individual and population level in the most challenging condition (i.e., at the −10 dB SNR).

### Reduction of amplitude modulation cues explains changes in neuronal discrimination

Tone vocoding degraded the spectro-temporal structure (i.e., it reduced both AM and FM cues) of original vocalizations in a deterministic way. Masking noise produced spectro-temporal degradations of the same nature, but it also introduced stochastic fluctuations in AM power. For each experimental condition (vocoding and noise), degradations of the spectro-temporal structure of vocalizations were quantified through the computation of AM and FM spectra at the output of a simulated cochlear filterbank. Not surprisingly, figure 1d (left panel) shows that tone vocoding corrupted drastically FM cues in each vocoding condition, and that this effect was already substantial in the 38-band vocoding condition. Importantly, this degradation was much stronger in vocoding conditions than in masking noise conditions (Fig. 1d). Figure 1e reveals that both tone vocoding and masking noise also attenuated the AM cues conveyed by vocalizations. There were only small AM degradations in the 38-band vocoding and in +10 dB SNR conditions; whereas important AM degradations were observed for the 0 dB SNR and the −10 dB SNR conditions. As for tone vocoding, this finding is consistent with the conclusions of previous modelling studies demonstrating the interplay between so-called temporal-envelope and TFS cues at the output of cochlear filters in response to vocoded sounds^16, 22^. Figure 5 relates these degradations of FM (Fig. 5a) and AM (Fig. 5b) cues to neural discrimination (MI_Population_) in the five brain structures for each experimental condition. More precisely, for all adverse conditions, figure 5 shows the changes in MI_Population_ for each auditory structure as a function of the attenuation of FM and AM cues (computed from modulation spectra for modulation rates between 1 and 20 Hz). Figure 5a reveals that an important attenuation of FM components caused by the 20-band vocoder was not associated with significant changes in neural discrimination in each structure, whereas a comparable attenuation caused by noise at −10 dB SNR resulted in a large drop in neural discrimination in each structure. Even more pronounced FM degradations caused by the 10-band vocoder condition produced smaller changes in MI_Population_ than the −10 dB SNR condition. This suggests that neural discrimination of vocalizations in the presence of background noise is not determined by the distortion of FM cues. A different pattern of results is obtained when changes in MI_Population_ for each auditory structure are represented as a function of degradations in AM cues. First, in all structures other than the CN, MI_Population_ is barely affected as long as the reduction of the AM index (Δmodulation index) remains lower than 25%; beyond this limit, the MI_Population_ is markedly reduced (i.e., at −10 dB SNR). The straightforward conclusion is that the reduction of AM cues is a key factor controlling the decrease in MI_Population_ at the cortical and subcortical levels. Second, in the cochlear nucleus, the impact on the MI_Population_ is much larger in the noise conditions (at 10 dB SNR and 0 dB SNR) than in the vocoding conditions, suggesting that the alteration of AM cues is not the only parameter driving the MI_Population_ at the most peripheral level. Noise and vocoding have in common to reduce the slow AM cues and corrupt the FM cues of vocalizations, but only background noise introduces randomness, that is stochastic fluctuations in AM power. Therefore, in the cochlear nucleus, the most likely factor responsible for the additional decrease in MI_Population_ is noise stochasticity. Third, in the condition causing the most severe alterations of AM cues, namely at −10 dB SNR, the MI_Population_ in CNIC is less impacted than in the other structures. Consistent with previous studies^23,24^, this finding suggests that inferior colliculus neurons adapt to the noise statistics while responding to the acoustic cues distinguishing between the four target stimuli.

**Figure 5.**
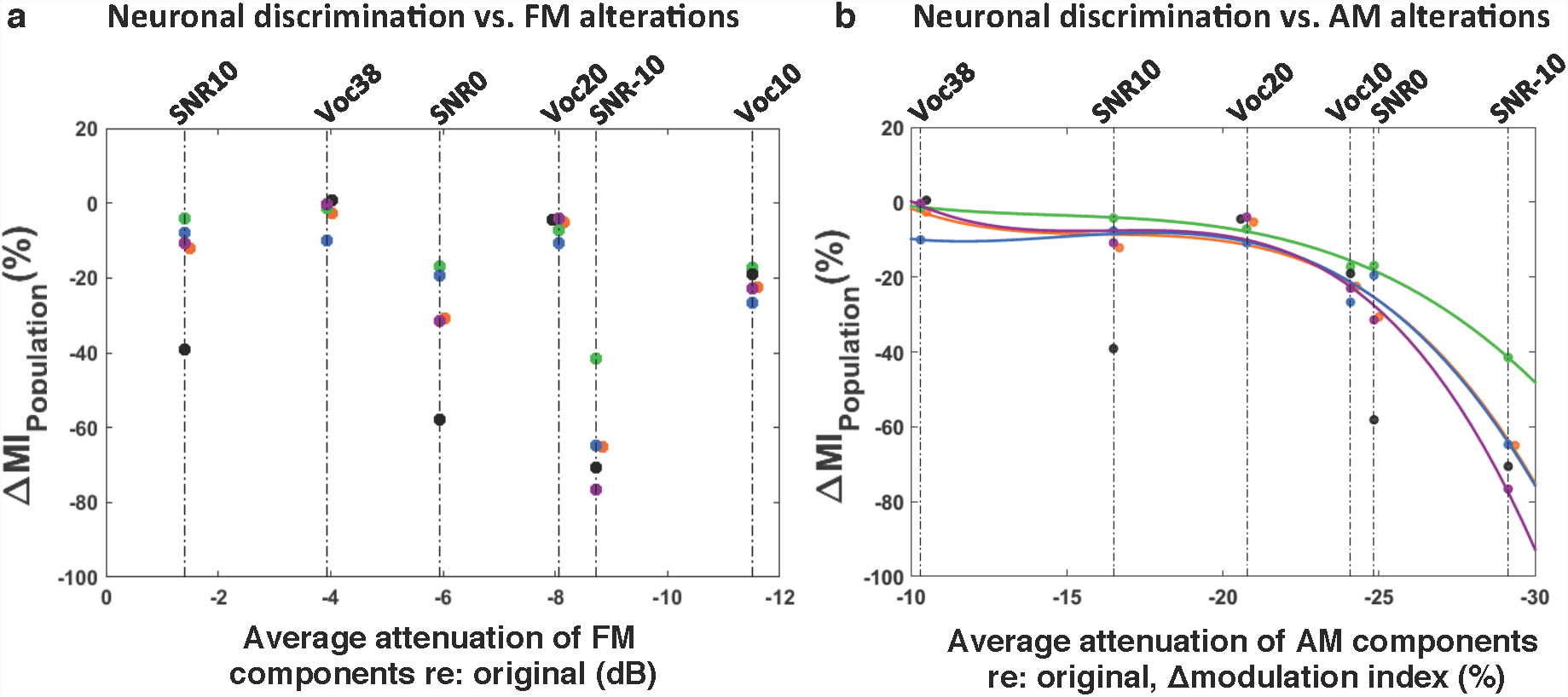
Reduction of AM cues determine the neuronal discriminative abilities at the subcortical and cortical levels. **a.** Percentage of ΔMI_Population_ as a function of degradation of FM components computed for each structure from means MI_Population_ or FM values obtained in all adverse conditions (noise and vocoded conditions) minus mean values in the original condition. Each dot represents neuronal data (ΔMI_Population_) in CN (*in black*), CNIC (*in green*), MGv (*in orange*), A1 (*in blue)* and VRB *(in purple).* **b.** Percentage of ΔMI_Population_ as a function of Δmodulation index computed for each structure from mean MI_Population_ or mean modulation-index values obtained in all adverse conditions and mean values in the original condition. Each dot represents neuronal data (ΔMI_Population_) in CN (*in black*), CNIC (*in green*), MGv (*in orange*), A1 (*in blue)* and VRB *(in purple)*. Polynomial curves fitting all acoustic conditions have been generated for each auditory structure (*color lines*) except for the cochlear nucleus. Note that equivalent AM degradations (Δmodulation index) induced equivalent decreases in MI_Population_ except for the CN. Note also that there is a limit of 25% of AM reduction beyond which the ΔMI_Population_ strongly decreases in cortical and subcortical structures.

## Discussion

Here, we demonstrate that the ability to discriminate between communication sounds is not increasing or decreasing monotonically along the auditory system: the neuronal discriminative abilities did strongly differ across auditory structures, and subcortical neurons in inferior colliculus and thalamus displayed higher discriminative abilities than cochlear nucleus and cortical neurons, both at the individual and population levels in each acoustic condition. Background noise markedly reduced the neuronal discriminative abilities in all auditory structures with larger effects in the cochlear nucleus. Amongst the three disruptive sub-effects of background noise identified in previous psychophysical investigations^12^, fidelity in the transmission of slow (< 20 Hz) amplitude modulation information proved to be the main factor determining the neural discrimination abilities in noise at the cortical and subcortical levels. The effects of randomness were found to be limited to the most peripheral structure of the central auditory system, namely the cochlear nucleus.

### The capacity to encode amplitude-modulation cues explains the better discrimination of the original stimuli by subcortical neurons

To the best of our knowledge, this is the first time that a direct comparison of neural responses to the same natural stimuli has been made along the ascending auditory system. We computed MI_Population_ in each structure from the cochlear nucleus to the secondary auditory cortex, and showed that on average subcortical populations discriminate the original vocalizations better than cortical populations. A much larger number of neurons exhibited high values of MI in the subcortical structures whatever the temporal precision considered (see supplementary figure 2); as a consequence, the growth of MI_Population_ as a function of the number of recordings included in the population increased more rapidly in the subcortical structures (see supplementary figure 3). The higher temporal reliability of subcortical neurons (higher CorrCoef values) probably allows them to follow more precisely the stimulus temporal envelope and encode more accurately the between-stimuli differences, both at the individual and population level.

To a large extent, our results corroborate those of Chechick et al. (2006)^25^ as we provide evidence that the neuronal discriminative abilities between communication sounds is higher in subcortical than in cortical structures. These authors showed that the MGB and AI responses contain 2-to-4 fold less information than the responses of IC neurons. Here, the neuronal discriminative ability of the ventral division of the auditory thalamus (MGv) was closer than the ones displayed by the other subcortical structures. A potential explanation is that Chechick et al. (2006)^25^ recorded from all divisions of the auditory thalamus, including the medial and dorsal divisions of the auditory thalamus, whereas our thalamic recordings were limited to the MGv and exhibited tonic responses to vocalizations similar to those observed in the CNIC and in the CN (see Fig. 2a and 3a). The stimulus sets also differ, as we used four utterances of the same category (the Whistle, an alarm call)^26^, whereas Chechick et al. (2006)^25^ used three birds chirps and variants of these stimuli such as the stimuli’s echoes or the background noises (15 stimuli in total), leading potentially to an easier classification between groups of stimuli compared to the present situation.

In our study, the original stimuli clearly differed in terms of AM patterns (i.e., their so-called “temporal envelope”) and, as a consequence, the most efficient way to discriminate them is probably to follow the time course of AM cues. It is well known that when progressing from the lower to the upper stages of the auditory system, the neurons’ ability to follow AM changes considerably^27,28^. Brainstem neurons phase-lock on the sounds’ AM pattern for AM rates up to hundreds of Hertz^29,30^, whereas thalamic neurons do so for a few tens of Hertz only^31,32^ and cortical neurons for even lower rates^33-35^. As a consequence, subcortical neurons are able to follow faster changes in the AM patterns of the original vocalizations, in addition to being more sensitive to spectro-temporal details. This likely explains why subcortical neurons better discriminate the original stimuli both at the individual and population level.

### Alterations of the slowest amplitude modulation cues explain changes in cortical and subcortical discrimination

Our main hypothesis is that the remarkable resistance of neural responses to the 0 dB SNR and the 10-band vocoded conditions results from the fact that AM cues were still sufficiently preserved. Previous studies using vocoded vocalizations reported little response changes at both the cortical and subcortical levels. In several species, cortical responses were not drastically reduced even when the number of frequency bands was reduced to two (marsomet^36^, gerbil^37^, rat^38^, guinea pig^39^). Most of the studies describing the effects of background noise on neuronal responses have been performed at the AI level, and many of them have pointed out the relationships between the noise impact on the cortical and behavioral discrimination performance. For example, in bird field L (homologous to AI), neuronal responses to song motifs were strongly reduced by three types of masking noises, and the neural discriminative ability was progressively reduced when the SNR decreased, in parallel with the behavioral performance^5^. Our VRB results are reminiscent of those obtained in the bird homologue of a secondary area (area NCM) where feed-forward inhibition, which potentially contributes to reduce the evoked discharges of pyramidal cells, allows the emergence of invariant neural representations of target songs in noise conditions^10^.

In mammals, the discriminative abilities of AI responses to speech sounds presented in quiet or against background noise closely match behavioral performance^40^. As in here, Shetake et al. (2011)^40^ did not find significant reduction in neural discrimination using an index of neuronal population performance (similar to MI) at a +12-dB SNR. Recent results revealed that responses of cortical neurons to calls could be classified in four classes-named robust, balanced, insensitive and brittle-when these calls were embedded in broadband white or babble noises^8^. In fact, the results of Bar-Yosef and Nelken (2007)^41^ in the cat primary auditory cortex have already shown that some neurons are more sensitive to the noise background than to the actual target stimulus. Here, we observed that the MI_Individual_ and MI_Population_ were only significantly reduced at a SNR of 0 dB, which indicates that on average, AI neurons were quite resistant to noise. This resistance is even higher in VRB where MI_Individual_ did not significant decrease and where MI_Population_ significantly decreased only at a SNR of −10 dB. In the only study performed at the subcortical level, responses of IC neurons were found to be resistant to drastic spectral degradations^42^. Here, we show that both at the individual and population level, the temporal reliability and discriminative ability of neurons in three subcortical auditory structures (CN, CNIC and MGv) are slightly but significantly reduced in the 10-band vocoded condition.

If our main hypothesis is valid, in situations where AM cues are attenuated either by vocoding or by noise, the neuronal discrimination based upon AM cues should be largely reduced. We showed that, in both conditions, the discriminative abilities decreased in each auditory structure when the AM index (Δmodulation index) was degraded by more than 25%. Therefore, these results indicate that accurate representation of slow AM cues is necessary at each level of the auditory system to discriminate efficiently communication sounds. More importantly, it appears that there is a limit to AM degradation beyond which the discriminative abilities decrease at the cortical and subcortical level. Thus, whatever the acoustic distortion used to degrade the amplitude modulations, equivalent AM alterations reduced the neuronal discrimination abilities to a similar extent. This is exactly what we observed for the Voc10 and 0 dB SNR conditions where degradations in the AM spectrum were comparable and produced the same decrease in neuronal discrimination in each structure, except in the cochlear nucleus. In this structure, discriminative abilities were more sensitive to noise than to vocoding, suggesting that the stochastic fluctuations introduced by noise impact the responses in cochlear nucleus, but not in the upper stages of the auditory system.

Only one previous study directly compared the impact of vocoding and masking noise on cortical responses to vocalizations^36^. This study shares several characteristics with our study. First, auditory cortex neurons were found to be robust to spectral degradations since there was little change in evoked firing rate, even in response to 2-band vocoded vocalizations. Second, broadband white noise reduced neuronal responses at 0 dB SNR. Third, temporal-envelope degradations strongly reduced the evoked firing rate and the neural synchronization to the vocalization envelope. Importantly, bandpass filtering the vocalizations between 2-30 Hz did not reduce firing rate and neural synchronization to the vocalization envelope. This is in total agreement with our results: when the AM index (Δmodulation index)-computed between 1 and 20 Hz – revealed modest AM alterations, there was little effect on the neuronal discrimination, but when the AM alterations were larger than 25%, the neuronal responses and neuronal discrimination were reduced (Fig. 5b). Thus, our results confirm that at the cortical level, the key factors constraining auditory discrimination are the slowest (< 20 Hz) AM cues and, importantly, extend this conclusion to subcortical structures.

Direct comparison between the consequences of acoustic degradations in different auditory structures using the same set of stimuli, anaesthetic agent and methods to quantify neural discrimination is the more straightforward way for dissecting where invariant representations are generated. When measuring how different levels of noise alter neuronal coding in the auditory system, it was found that the neural representation of natural sounds becomes progressively independent of the level of background noise from the auditory nerve till the IC and AI^23^. It was proposed that at the population level, this tolerance to background noise results from an adaptation to the noise statistics, which is much more pronounced at the cortical than at the subcortical level^23^. In agreement with this study, we found that populations of cortical neurons (AI and VRB) were more resistant to noise than subcortical ones. However, we did not observe a monotonic evolution of resistance to noise in the auditory system: at the subcortical level, the discrimination abilities of CN neuronal populations drastically dropped for a +10 dB SNR and those of thalamic ones largely decreased at a −10 dB SNR, whereas populations of CNIC cells maintained relatively good discriminative abilities, suggesting that they were the more resistant to noise, even in the most adverse conditions. In the IC, previous work showed that background noise changes the shape of the temporal modulation transfer function of individual neurons from bandpass to lowpass^24^. The CNIC is a massive hub receiving probably the highest diversity of inhibitory and excitatory inputs^43,44^ and potentially the large diversity of these inputs allows this structure to extract crucial temporal information about the stimulus’ temporal envelope, even at relatively low SNR.

### General conclusions

The present study led to two major findings with regard to the main factors influencing auditory processing in noise, and the respective contributions of auditory structures to robust sound coding in noise.

Comparison of neural data collected in response to noisy versus vocoded vocalizations clarified a long-lasting debate^11;12,16,45^: from a neural perspective, the *main* effect of (notionally) steady background noise on complex-sound discrimination corresponds to the attenuation of the gross AM (i.e., “temporal envelope”) cues conveyed by sounds. Corruption of FM cues and introduction of stochastic fluctuations in AM power have little influence if any on neural discrimination in noise (with the noticeable exception of the cochlear nucleus showing strong sensitivity to stochasticity). This is in accordance with objective measures currently used by audio engineers to predict the intelligibility and perceived quality of speech masked by noise^13, 45^.

Inconsistent with our initial expectations, the ability of auditory neurons to discriminate between communication sounds masked by external noise neither increased nor decreased along the auditory pathway from the first auditory relay up to the primary and secondary cortical areas, but culminated at the collicular and thalamic levels. In humans, speech sounds (such as phonemes) showing similar acoustic properties trigger similar responses and are represented as a single category in the superior temporal gyrus^4^. Here, the use of vocalizations belonging to the same category of the communication repertoire of guinea pigs, i.e. “whistles”, may explain both the relatively poor discriminative abilities of cortical neurons compared to subcortical ones and the robustness of cortical responses to vocoding and background noise. In these two challenging situations, our results reveal that cortical neurons resist more than the other auditory structures, potentially because cortical neurons do not code for the spectro-temporal details of the stimuli but rather respond to more abstract stimulus properties^21^ carried by the gross spectro-temporal envelope patterns.

In a recent study, auditory cortex responses collected in behaving ferrets were found to be sufficiently robust to preserve vowel identity across a large range of acoustic transformations, such as changes in fundamental frequency, sound location or level^9^. It is notable that earlier studies from the same laboratory performed in anaesthetized conditions^46,47^ have reached very similar conclusions for vowels varying in fundamental frequency and virtual acoustic location, indicating that the general principles allowing neuronal discrimination are observable across anesthetized and behavioral states.

Our results extend these findings by showing that downstream from AI, neurons in a secondary auditory area (VRB) are even more resistant to spectral degradations than in AI. This is in line with the results of Carruthers et al. (2015)^7^ showing that in the secondary auditory cortex (the SRAF area), neuronal populations code invariant representations of conspecific vocalizations despite important spectro-temporal degradations. As already proposed^21^ by Chechick and Nelken (2012)^21^, auditory cortex neurons extract abstract auditory entities rather than detailed spectro-temporal features. This suggests that even when the four vocalizations were vocoded, the mere fact that gross spectro-temporal envelope cues were preserved in a limited number of frequency bands was sufficient for auditory cortex neurons to classify these stimuli as belonging to the alarm-call category.

Neurons in subcortical structures were able to discriminate better the stimuli than the neurons in cortical areas when temporal envelope cues were degraded even in the most severe condition (>25% of alterations in the −10 dB SNR condition). Indeed, the most accurate representations of target stimuli (and of their differences) were found at the collicular and thalamic levels, not at the cortical level. In each condition, the identification of an auditory object necessarily involves both subcortical and cortical processing. We therefore suggest that in challenging conditions, cortical representations co-exist with more detailed representations of the stimuli in one, or several, subcortical structures. In fact, in the most adverse noise condition (at a −10 dB SNR) where temporal envelope cues were strongly degraded, CNIC neurons still exhibited discrimination abilities whereas the other subcortical and cortical neurons did not (a value of 1 for the MI_Population_ indicates that 2 out of 4 stimuli could still be discriminated). Further studies are required to determine to what extent these incomplete subcortical representations influence auditory abilities in animals and humans.

## Materials and Methods

### Subjects

These experiments were performed under the national license A-91-557 (project 2014-25, authorization 05202.02) and using the procedures N° 32-2011 and 34-2012 validated by the Ethic committee N°59 (CEEA Paris Centre et Sud). All surgical procedures were performed in accordance with the guidelines established by the European Communities Council Directive (2010/63/EU Council Directive Decree).

Extracellular recordings were obtained from 47 adult pigmented guinea pigs (aged 3 to 16 months) at five different levels of the auditory system: the cochlear nucleus (CN), the inferior colliculus (IC), the medial geniculate body (MGB), the primary (AI) and secondary (area VRB) auditory cortex. Animals weighting from 515 to 1100 g (mean 856 g) came from our own colony housed in a humidity (50-55%) and temperature (22-24°C)-controlled facility on a 12 h/12 h light/dark cycle (light on at 7:30 A.M.) with free access to food and water.

Two to three days before each experiment, the animal’s pure-tone audiogram was determined by testing auditory brainstem responses (ABR) under isoflurane anaesthesia (2.5 %) as described in Gourévitch et al. (2009)^48^. The ABR was obtained by differential recordings between two subdermal electrodes (SC25-NeuroService) placed at the vertex and behind the mastoid bone. A software (RTLab, Echodia, Clermont-Ferrand, France) allowed averaging 500 responses during the presentation of nine pure-tone frequencies (between 0.5 and 32 kHz) delivered by a speaker (Knowles Electronics) placed in the animal right ear. The auditory threshold of each ABR was the lowest intensity where a small ABR wave can still be detected (usually wave III). For each frequency, the threshold was determined by gradually decreasing the sound intensity (from 80 dB down to −10 dB SPL). All animals used in this study had normal pure-tone audiograms^39,48,49^.

### Surgical procedures

All animals were anesthetized by an initial injection of urethane (1.2 g/kg, i.p.) supplemented by additional doses of urethane (0.5 g/kg, i.p.) when reflex movements were observed after pinching the hind paw (usually 2-4 times during the recording session). A single dose of atropine sulphate (0.06mg/kg, s.c.) was given to reduce bronchial secretions and a small dose of buprenorphine was administrated (0.05mg/kg, s.c.) as urethane has no antalgic properties. After placing the animal in a stereotaxic frame, a craniotomy was performed and a local anesthetic (Xylocain 2%) was liberally injected in the wound.

For auditory cortex recordings (area A1 and VRB), a craniotomy was performed above the left temporal cortex. The opening was 8 mm wide starting at the intersection point between parietal and temporal bones and 8-10 mm height. The dura above the auditory cortex was removed under binocular control and the cerebrospinal fluid was drained through the cisterna to prevent the occurrence of oedema.

For the recordings in MGB, a craniotomy was performed the most posterior part of the MGB (8mm posterior to Bregma) to reach the left auditory thalamus at a location where the MGB is mainly composed of its ventral, tonotopic, part^50-52^.

For IC recordings, a craniotomy was performed above the IC and portions of the cortex were aspirated to expose the surface of the left IC. For CN recordings, after opening the skull above the right cerebellum, portions of the cerebellum were aspirated to expose the surface of the right CN^53^.

After all surgery, a pedestal in dental acrylic cement was built to allow an atraumatic fixation of the animal’s head during the recording session. The stereotaxic frame supporting the animal was placed in a sound-attenuating chamber (IAC, model AC1). At the end of the recording session, a lethal dose of Exagon (pentobarbital >200 mg/kg, i.p.) was administered to the animal.

### Recording procedures

Data were from multi-unit recordings collected in 5 auditory structures, the non-primary cortical area VRB, the primary cortical area A1, the medial geniculate body (MGB), the inferior colliculus (IC) and the cochlear nucleus (CN). Cortical extracellular recordings were obtained from arrays of 16 tungsten electrodes (ø: 33 µm, <1 MΩ) composed of two rows of 8 electrodes separated by 1000 µm (350 µm between electrodes of the same row). A silver wire, used as ground, was inserted between the temporal bone and the dura matter on the contralateral side. The location of the primary auditory cortex was estimated based on the pattern of vasculature observed in previous studies^54-57^. The non-primary cortical area VRB was located ventral to A1 and distinguished because of its long latencies to pure tones^58,59^. For each experiment, the position of the electrode array was set in such a way that the two rows of eight electrodes sample neurons responding from low to high frequency when progressing in the rostro-caudal direction [see examples of tonotopic gradients recorded with such arrays in figure 1 of Gaucher et al. (2012)^60^ and in figure 6A of Occelli et al. (2016)^61^.

All the remaining extracellular recordings (in MGB, IC and CN) were obtained using 16 channel multi-electrode arrays (NeuroNexus) composed of one shank (10 mm) of 16 electrodes spaced by 110 µm and with conductive site areas of 177µm^2^. The electrodes were advanced vertically (for MGB and IC) or with a 40° angle (for CN) until evoked responses to pure tones could be detected on at least 10 electrodes.

All thalamic recordings were from the ventral part of MGB (see above surgical procedures) and all displayed latencies < 9ms. At the collicular level, we distinguished the lemniscal and non-lemniscal divisions of IC based on depth and on the latencies of pure tone responses. We excluded the most superficial recordings (until a depth of 1500µm) and those exhibiting latency >= 20ms in an attempt to select recordings from the central nucleus of IC (CNIC). At the level of the cochlear nucleus, the recordings were collected from both the dorsal and ventral divisions.

The raw signal was amplified 10,000 times (TDT Medusa). It was then processed by an RX5 multichannel data acquisition system (TDT). The signal collected from each electrode was filtered (610-10000 Hz) to extract multi-unit activity (MUA). The trigger level was set for each electrode to select the largest action potentials from the signal. On-line and off-line examination of the waveforms suggests that the MUA collected here was made of action potentials generated by 2 to 6 neurons in the vicinity of the electrode.

### Acoustic stimuli

Acoustic stimuli were generated using MatLab, transferred to a RP2.1-based sound delivery system (TDT) and sent to a Fostex speaker (FE87E). The speaker was placed at 2 cm from the guinea pig’s right ear, a distance at which the speaker produced a flat spectrum (± 3 dB) between 140 Hz and 36 kHz. Calibration of the speaker was made using noise and pure tones recorded by a Bruel & Kjaer microphone 4133 coupled to a preamplifier B&K 2169 and a digital recorder Marantz PMD671.

Spectro-temporal receptive fields (STRFs) were first determined using 97 or 129 pure-tones frequencies scaled with a gamma function, covering six (0.14-9 kHz or 0.28-18 kHz or 0.56-36 kHz) or eight (0.14-36 kHz) octaves respectively, and presented at 75 dB SPL. At a given level, each frequency was repeated eight times at a rate of 2.35 Hz in pseudorandom order.

The duration of these tones over half-peak amplitude was 15 ms and the total duration of the tone was 50 ms, so there was no overlap between tones.

A set of four conspecific vocalizations was used to assess the neuronal responses to communication sounds. These vocalizations were recorded from animals of our colony. Pairs of animals were placed in the acoustic chamber and their vocalizations were recorded by a Bruel & Kjaer microphone 4133 coupled to a preamplifier B&K 2169 and a digital recorder Marantz PMD671. A large set of whistle calls was loaded in the Audition software (Adobe Audition 3) and four representative examples of whistle were selected. As shown in figure 1a (lower panels), despite the fact the maximal energy of the four selected whistle was in the same frequency range (typically between 4 and 26 kHz), these calls displayed slight differences in their spectrograms. In addition, their temporal (amplitude) envelopes clearly differed as shown by their waveforms (Fig. 1a, upper panels). The four selected whistles were processed by three tone vocoders^62,63^. In the following figures, the unprocessed whistles will be referred to as the original versions, and the vocoded versions as Voc38 (Voc20, Voc10 respectively) for the 38-band (20-band, 10-band, respectively) vocoded whistles. In contrast to previous studies that used noise-excited vocoders^36-38^, a tone vocoder was used here, because noise vocoders were found to introduce random (i.e., non-informative) intrinsic temporal-envelope fluctuations distorting the crucial spectro-temporal modulation features of communication sounds^16, 22,64^.

Figure 1b displays the spectrograms of the 38-band vocoded (first row), the 20-band vocoded (second row) and the 10-band vocoded (third row) of the four whistles. The three vocoders differed only in terms of the number of frequency bands (i.e., analysis filters) used to decompose the whistles (38, 20 or 10 bands). The 38-band vocoding process is briefly described below, but the same principles apply to the 20-band or the 10-band vocoders. Each digitized signal was passed through a bank of 38 fourth-order Gammatone filters^65^ with center frequencies uniformly spaced along a guinea-pig adapted ERB (Equivalent Rectangular Bandwidth) scale^66^ ranging from 50 to 35505 Hz. In each frequency band, the temporal envelope was extracted using full-wave rectification and lowpass filtering at 64 Hz with a zero-phase, sixth-order Butterworth filter. The resulting envelopes were used to amplitude modulate sine-wave carriers with frequencies at the center frequency of the Gammatone filters, and with random starting phase. Impulse responses were peak-aligned for the envelope (using a group delay of 16 ms) and the acoustic temporal fine structure across frequency channels^67^. The modulated signals were finally weighted and summed over the 38 frequency bands. The weighting compensated for imperfect superposition of the bands’ impulse responses at the desired group delay. The weights were optimized numerically to achieve a flat frequency response. Amplitude-modulation (AM) spectra were computed for the original and vocoded versions of each vocalization and the averaged modulation spectra are displayed in figure 1d (left panel). AM spectra were computed^17^ by decomposing each sound using a bank of 50 (spanning the range 0.1-50 kHz), 1 ERB-wide Gammatone filters and then analyzing the temporal envelope in each frequency band through a bank of AM filters (1/3-octave wide first-order Butterworth bandpass filters overlapping at −3 dB, with center frequencies between 0.1 Hz and 410 Hz). The root-mean-square amplitude of the filtered output was multiplied by a factor of 1.414. For each AM filter, a modulation index was calculated by dividing the output by the mean amplitude of the AM component for the vocalization sample in a given Gammatone filter. Finally, the 50 AM spectra were averaged to generate a single AM spectrum per stimulus. All vocalizations were presented at 75dB SPL.

The four whistles were also presented in a frozen stationary background noise ranging from 100-30000 Hz. The first three rows of figure 1c display the spectrograms of the four whistles in the stationary noise with a SNR of +10 dB SPL, 0 dB SPL, −10 dB SPL and the last row shows the masking noise only. The alterations of the AM spectra produced by masking noise on the original vocalizations are displayed in figure 1d (right panel). To quantify the alterations of the temporal envelope induced either by the noise addition or by the vocoding, we calculated the mean percentage of variation of the modulation index (Δmodulation index, computed from 1-20 Hz) between each acoustic condition (noise or vocoded condition) and the original condition (Fig. 5b).

### Experimental protocol

As inserting an array of 16 electrodes in a brain structure almost systematically induces a deformation of this structure, a 30-minutes recovering time lapse was allowed for the structure to return to its initial shape, then the array was slowly lowered. Tests based on measures of spectro-temporal receptive fields (STRFs) were used to assess the quality of our recordings and to adjust electrodes’ depth. For auditory cortex recordings (AI and VRB), the recording depth was 500-1000 µm, which corresponds to layer III and the upper part of layer IV according to Wallace and Palmer (2008)^68^. For thalamic recordings, the NeuroNexus probe was lowered of about 7mm below pia before the first responses to pure tones were detected.

When a clear frequency tuning was obtained for at least 10 of the 16 electrodes, the stability of the tuning was assessed: we required that the recorded neurons displayed at least three successive similar STRFs (each lasting 6 minutes) before starting the protocol. When the stability was satisfactory, the protocol was started by presenting the acoustic stimuli in the following order: We first presented the 4 whistles at 75 dB SPL in their natural versions, followed by the vocoded versions with 38, 20 and 10 bands. The same set of original whistles was then presented in the stationary noise presented at 65, 75 and 85 dB SPL. In each case, each vocalization was repeated 20 times. Presentation of this entire stimulus set lasted 45 minutes. The protocol was re-started either after moving the electrode arrays on the cortical map or after lowering the electrode at least by 300 µm for subcortical structures.

### Data analysis

#### Quantification of responses to pure tones

The STRFs derived from MUA were obtained by constructing post-stimulus time histograms for each frequency with 1 ms time bins. The firing rate evoked by each frequency was quantified by summing all the action potentials from the tone onset up to 100 ms after this onset. Thus, STRFs are matrices of 100 bins in abscissa (time) multiplied by 97 or 129 bins in ordinate (frequency). All STRFs were smoothed with a uniform 5×5 bin window.

For each STRF, the Best Frequency (BF) was defined as the frequency at which the highest firing rate was recorded. Peaks of significant response were automatically identified using the following procedure: A positive peak in the MU-based STRF was defined as a contour of firing rate above the average level of the baseline activity (estimated from the ten first milliseconds of STRFs at all intensity levels) plus six times the standard deviation of the baseline activity.

#### Quantification of responses evoked by vocalizations

The responses to vocalizations were quantified using two parameters: (i) The firing rate of the evoked response, which corresponds to the total number of action potentials occurring during the presentation of the stimulus minus spontaneous activity; (ii) the spike-timing reliability coefficient (CorrCoef) which quantifies the trial-to-trial reliability of the response. This index was computed for each vocalization: it corresponds to the normalized covariance between each pair of spike trains recorded at presentation of this vocalization and was calculated as follows:

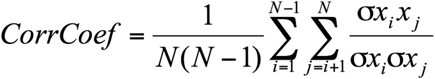

where N is the number of trials and *σx_i_x_j_* is the normalized covariance at zero lag between spike trains x_i_ and x_j_ where i and j are the trial numbers. Spike trains x_i_ and x_j_ were previously convolved with a 10-msec width Gaussian window. Based upon computer simulations, we have previously shown that this CorrCoef index is not a function of the neurons’ firing rate^69^. We have computed the CorrCoef index with a Gaussian window ranging from 1 to 50 ms to determine if the selection of a particular value for the Gaussian window influences the difference in CorrCoef mean values obtained in the different auditory structures. As a general rule, the largest the Gaussian window, the largest the CorrCoef value whatever the structure was. Based upon the responses to the original vocalizations, supplementary Figure S1A shows that the relative ranking between auditory structures remained unchanged whatever the size of the Gaussian window was. Therefore, we kept the value of 10 ms for the Gaussian window (dashed line in figure S1) as it was used in several previous studies^39, 69-71^.

#### Quantification of mutual information from the responses to vocalizations

The method developed by Schnupp et al. (2006)^72^ was used to quantify the amount of information (Shannon 1948) contained in the responses to vocalizations obtained with natural vocoded and noise stimuli. This method allows quantifying how well the vocalization’s identity can be inferred from neuronal responses. Here, “neuronal responses” refer either to (i) the spike trains obtained from a small group of neurons below one electrode (for the computation of the individual Mutual Information, MI_Individual_), or to (ii) a concatenation of spike trains simultaneously recorded under several electrodes (for the computation of the population, MI_Population_). In both cases, the following computation steps were the same. Neuronal responses were represented using different time scales ranging from the duration of the whole response (firing rate) to a 1-ms precision (precise temporal patterns), which allows analyzing how much the spike timing contributes to the information. As this method is exhaustively described in Schnupp et al. (2006)^72^ and in Gaucher et al. (2013a)^69^, we only present below the main principles.

The method relies on a pattern-recognition algorithm that is designed to “guess which stimulus evoked a particular response pattern”^72^ by going through the following steps: From all the responses of a cortical site to the different stimuli, a single response (test pattern) is extracted and represented as a PSTH with a given bin size (different sizes were considered as discussed further below). Then, a mean response pattern is computed from the remaining responses (training set) for each stimulus class. The test pattern is then assigned to the stimulus class of the closest mean response pattern. This operation is repeated for all the responses, generating a confusion matrix where each response is assigned to a given stimulus class. From this confusion matrix, the Mutual Information (MI) is given by Shannon’s formula:

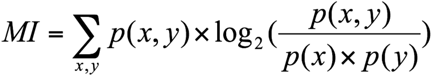

where x and y are the rows and columns of the confusion matrix, or in other words, the values taken by the random variables “presented stimulus class” and “assigned stimulus class”.

In our case, we used responses to the 4 whistles and selected the first 264 ms of these responses to work on spike trains of exactly the same duration (the shortest whistle being 280msec long). In a scenario where the responses do not carry information, the assignments of each response to a mean response pattern is equivalent to chance level (here 0.25 because we used 4 different stimuli and each stimulus was presented the same number of times) and the MI would be close to zero. In the opposite case, when responses are very different between stimulus classes and very similar within a stimulus class, the confusion matrix would be diagonal and the mutual information would tend to log_2_(4) =2 bits.

This algorithm was applied with different bin sizes ranging from 1 to 280 ms. Supplementary figure S1B presents the evolution of MI as a function of temporal precision ranging from 1 to 40ms with the responses to the original vocalizations. At the cortical level, we previously showed that an optimal bin size for obtaining a maximal value of MI was on average 8ms^69,72^. However, it has never been demonstrated that the same bin size value was optimal at the thalamic and brainstem levels. In our experimental conditions, and with our set of acoustic stimuli, the 8-ms temporal precision was found to be optimal for all auditory structures (dashed line in supplementary Fig. 1b). The MI value obtained for a temporal precision of 8ms was subsequently used in our analysis.

The MI estimates are subject to non-negligible positive sampling biases. Therefore, as in Schnupp et al. (2006)^72^, we estimated the expected size of this bias by calculating MI values for “shuffled” data, in which the response patterns were randomly reassigned to stimulus classes. The shuffling was repeated 100 times, resulting in 100 MI estimates of the bias (MI_bias_). These MI_bias_ estimates are then used as estimators for the computation of the statistical significance of the MI estimate for the real (unshuffled) datasets: the real estimate is considered as significant if its value is statistically different from the distribution of MI_bias_ shuffled estimates. Significant MI estimates were computed for MI calculated from neuronal responses under one electrode.

The information carried by a group of recording was estimated by the population MI (MI_Population_), using the same method described above: responses of several simultaneous recordings were grouped together and considered as a single pattern. To assess the influence of the group size of simultaneous recordings on the information carried by that group (MI_Population_), the number of sites used for computing MI_Population_ varied from 2 to the maximal possible size (which is equal to 16 minus the non-responsive sites). As the number of possible combinations could be extremely large (C_n_^k^, where k is the group size and n the number of responsive sites in a recording session), a threshold was fixed to save computation time: when the number of possible combinations exceeded one hundred, 100 combinations were randomly chosen, and the mean of all combinations was taken as the MI_Population_ for this group size.

#### Statistics

To assess the significance of the multiple comparisons (Vocoding process: four levels; Masking noise conditions: three levels; Brain structure: five levels), we used an analysis of variance (ANOVA) for multiple factors to analyze the whole data set. Follow-up tests were corrected for multiple comparisons using Bonferroni corrections and were considered as significant if their p value was below 0.05. All data are presented as mean values ± standard error (s.e.m.).

---

**Supplementary figure 1.**
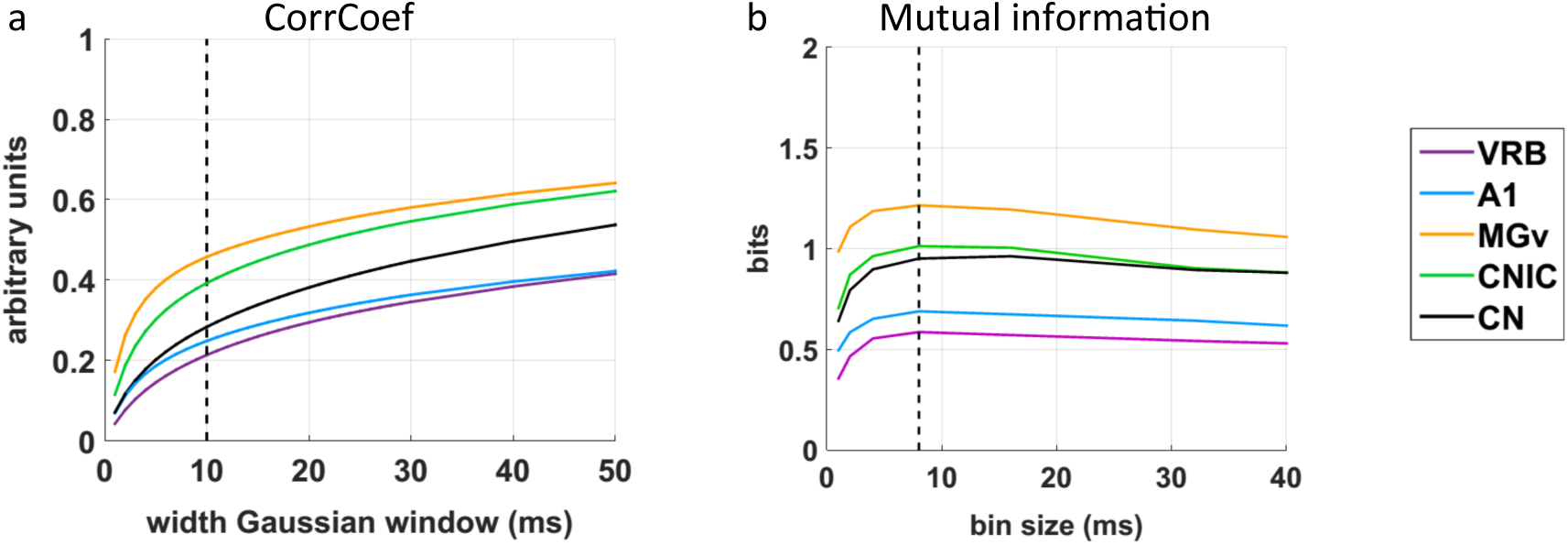
Evolution of the CorrCoef and MI mean values as a function of temporal precision in each structure. **a.** CorrCoef values were calculated from responses to original vocalisations with a Gaussian window varying in width from 1 to 50 ms in CN (*in black*), CNIC (*in green*), MGv (*in orange*), A1 (*in blue)* and VRB *(in purple)*. In our study, a 10-ms width Gaussian window (*dashed black line*) was selected for the data analysis in each structure. **b.** Mutual information (in *bits*) was calculated from neuronal responses to original vocalizations with a bin size varying from 1 to 40 ms in CN (*in black*), CNIC (*in green*), MGv (*in orange*), A1 (*in blue)* and VRB *(in purple)*. In this study, the value of 8 ms was selected for the data analysis because in each structure, the MI value was maximal (*dashed black line*).

**Supplementary figure 2.**
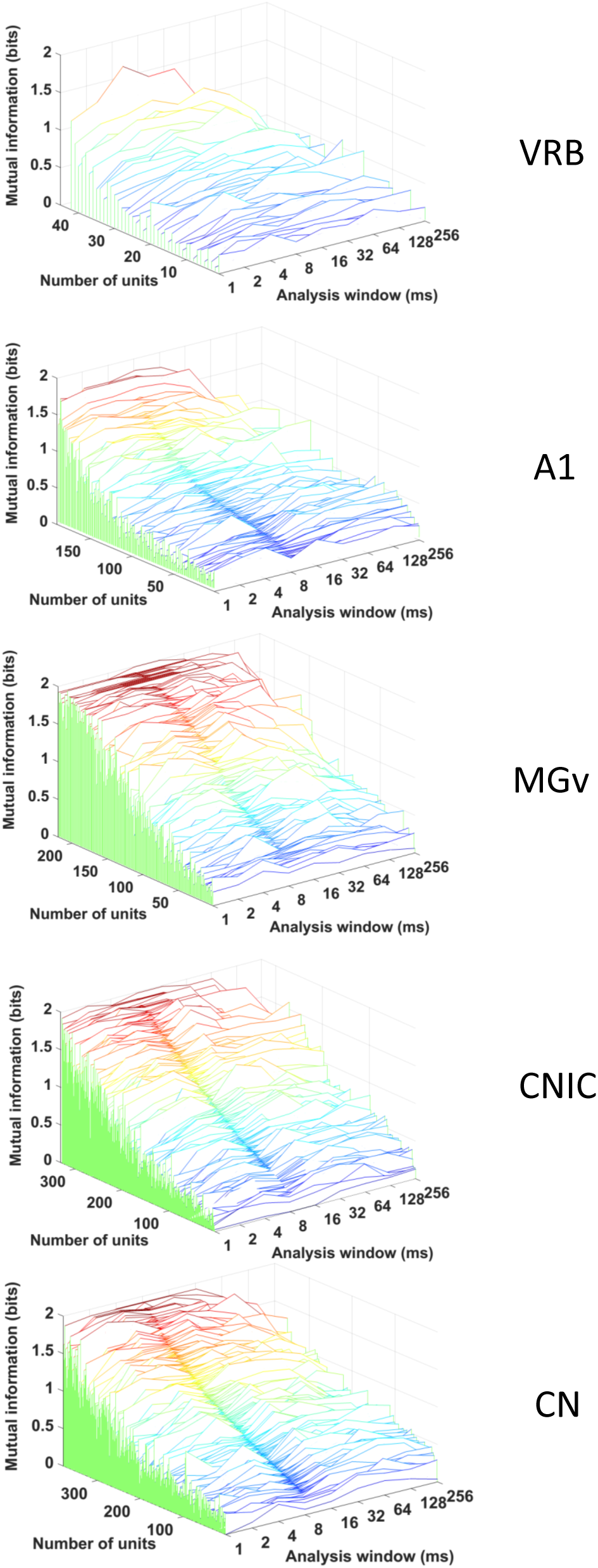
Large diversity of neuronal discrimination performance in response to original vocalizations in each auditory structure. Waterfall plots show the mutual information (MI, *bits*) as a function of temporal resolution (1 to 256 ms) for the selected recordings in CN, CNIC, MGv, A1 and VRB. In each structure, the units are ranked by the MI value obtained with a bin size of 8ms (*dark rainbow colors*). Note that there was a larger proportion of neurons with high values of MI (close from the maximal value of 2) in MGv, CNIC and CN (*red curves*) compared to a much lower proportion in the cortical areas AI and VRB.

**Supplementary figure 3.**
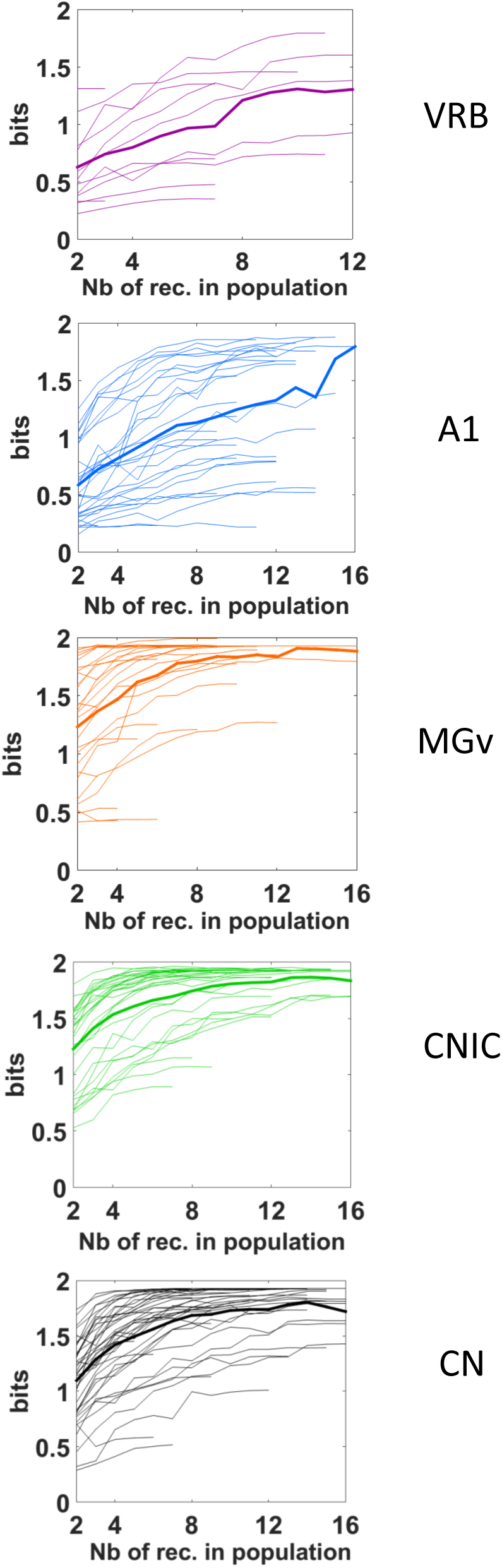
Population information quickly reaches high values with simultaneous recordings at the subcortical but not cortical level. For each auditory structure, each thin line represents a particular case of simultaneous recordings with a maximum number of electrodes varying from 2 to 12 or 16, and each thick line represents the mean value of MI_Population_ in CN (*in black*), CNIC (*in green*), MGv (*in orange*), A1 (*in blue)* and VRB *(in purple)*. Note that the mean MI_Population_ value quickly reaches high values close from the maximum value of 2 bits in the subcortical structures (CN, CNIC and MGv) compared to the two cortical areas (A1 and VRB).

**Supplementary figure 4.**
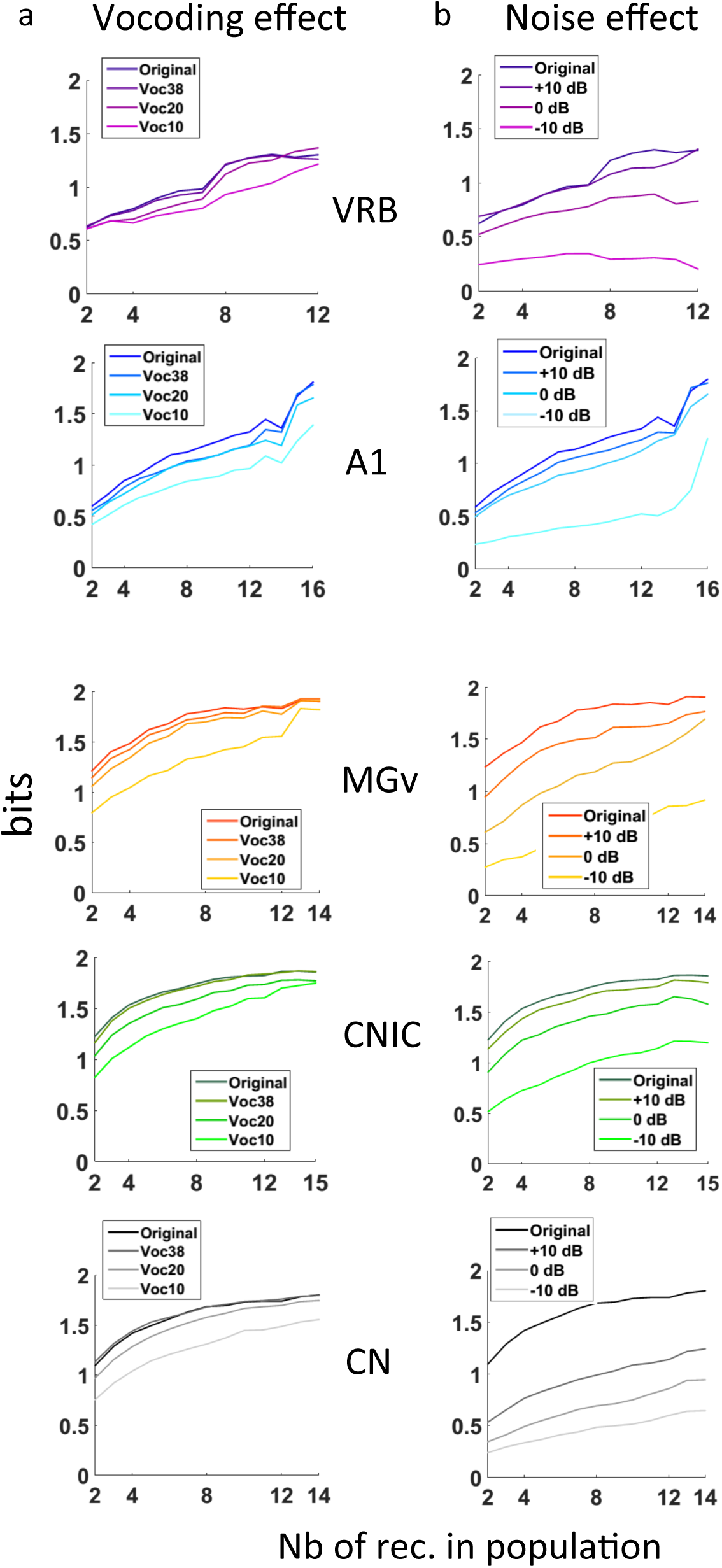
Effects of vocoding and noise on the MI_Population_ growth functions in each auditory structure. a. Vocoding effects. The graphics display the average growth functions of the MI_Population_ for each structure in each vocoding condition (indicated by a gradient colors). In each structure, the vocoding slightly reduced the MI_Population_ values. At the cortical level, the reduction induced by vocoding was similar at 38 and 20 bands, then a stronger reduction was observed at 10 bands. At the thalamic level, there was almost no change in the growth function of the MI_Population_ with 38 and 20 bands vocalizations, but there was a large decrease in MI_Population_ with the 10 bands vocoded stimuli. In the CNIC, there was almost no change in the growth function of the MI_Population_ with 38-bands vocalizations, and a progressive reduction of MI_Population_ with 20 and 10-bands vocalizations. A similar scenario was observed at the CN level. **b. Noise effects.** The graphics display the effects noise on the growth functions of the MI_Population_ for each structure and at each SNR. In general, background noise largely altered the growth functions of the MI_Population_ in each structure (but to a lesser extent in the CNIC). In the cortex, SNR affected the growth functions of the MI_Population_ : the lower the SNR, the lower the curves of the MI_Population_. In the MGv, stationary noise progressively lowered the curves of the MI_Population_. In the CNIC, stationary noise induced SNR-dependent reduction in the MI_Population_ values, the reduction being modest at a +10 and 0 dB SNR but more important at a-10 dB SNR. In the CN, stationary noise induced a stronger reduction of the MI_Population_ which was clearly a function of SNR.

**Supplementary figure 5.**
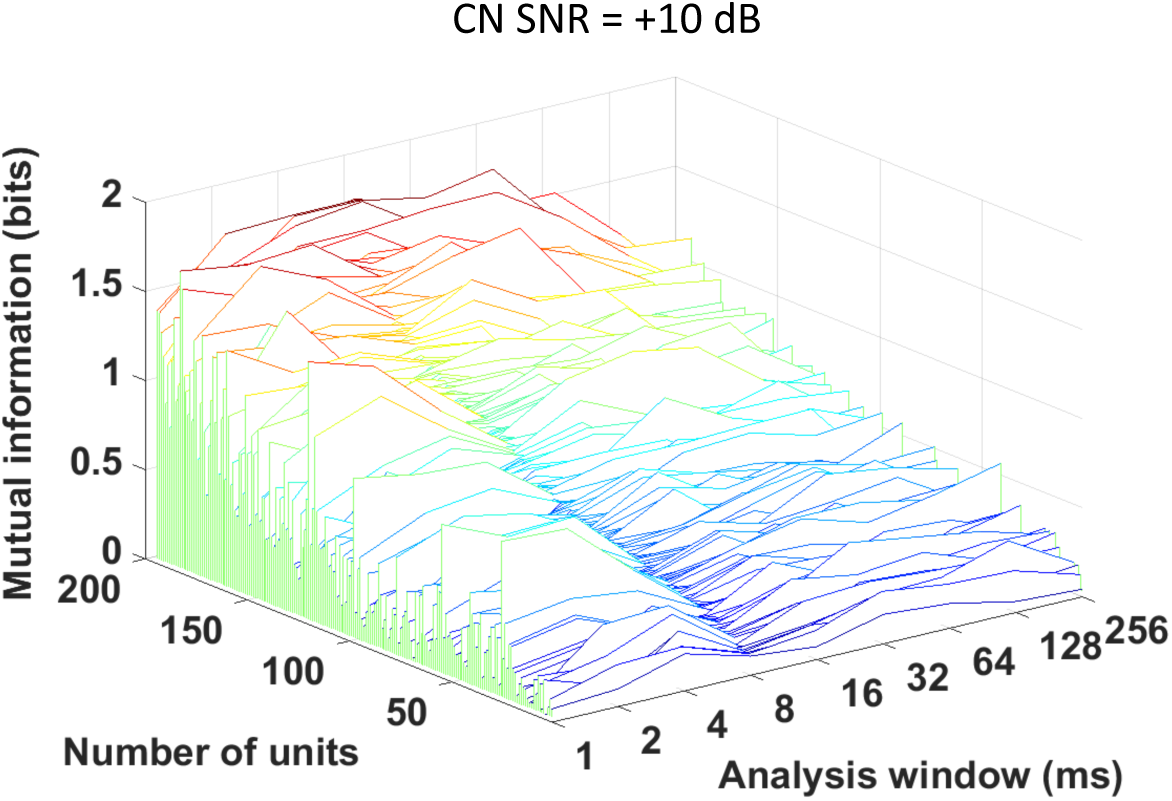
A subpopulation of CN neurons maintains good neuronal discrimination performance at a +10 dB SNR. Waterfall plots show the mutual information (MI_Individual_, bits) as a function of temporal resolution (1 to 256 ms) for the CN recordings at a +10 dB SNR. The recordings are ranked by the MI value obtained with a bin size of 8 ms (*dark rainbow colors*). Note that at this particular SNR, 20% of the CN recordings (n=39) maintained MI_Individual_ values above 1 bit, indicating that some CN neurons still send information about the vocalization identity at higher brainstem centers such as the CNIC.

**Supplementary Table 1.** Summary of the number of animals, number of selected recordings and STRFs quantifications in each structure.

|  | CN | Lemniscal pathway |  |  | Non-lemniscal pathway |
| --- | --- | --- | --- | --- | --- |
|  |  | CNIC | MGv | A1 | VRB |
| Number of animals | 10 | 11 | 10 | 11 | 5 |
| Number tested | 672 | 478 | 448 | 544 | 192 |
| STRF only | 560 | 421 | 285 | 455 | 126 |
| Selection criteria : STRF & response to at least one vocalization | 499 | 386 | 262 | 354 | 95 |
| **STRF quantifications** |  |  |  |  |  |
| BF range (kHz) : min-max | 0.18 - 18 | 0.34 - 36 | 0.33 - 33.01 | 0.14 - 36 | 0.67 - 36 |
| Mean bandwidth (octave) | 3.91 | 2.88 | 4.16 | 2.07 | 1.79 |
| Mean response duration (ms) | 26.83 | 35.37 | 17.31 | 43.73 | 44.83 |
| Response strength (spikes/sec) | 77.23 | 82.25 | 41.61 | 37.69 | 19.97 |

## Acknowledgments

CL and JME were supported by grants from the French Agence Nationale de la Recherche (ANR) (ANR-14-CE30-0019-01). CL was also supported by grants ANR-11-0001-02 PSL and ANR-10-LABX-0087. SS was supported by the Fondation pour la Recherche Médicale (FRM) grant number ECO20160736099.

We thank warmly Léo Varnet for help in analyzing the modulation spectra of the acoustic stimuli and Roger Mundry for detailed and relevant comments on statistical analyses. We thank Nihaad Paraouty for training us to cochlear-nucleus surgery. We also wish to thank Mélanie Dumont, Aurélie Bonilla and Céline Dubois for taking care of the guinea-pig colony. We warmly thank Dr. Dexter Irvine and Dr. Jonathan Fritz for helpful comments on an earlier draft of this paper.

